# Central Role of Cognitive Control Networks in Weight Loss During Voluntary Calorie Restriction

**DOI:** 10.1101/234641

**Authors:** Selin Neseliler, Wen Hu, Kevin Larcher, Maria Zacchia, Mahsa Dadar, Stephanie G. Scala, Marie Lamarche, Yashar Zeighami, Stephen C. Stotland, Maurice Larocque, Errol B. Marliss, Alain Dagher

## Abstract

Insufficient responses to hypocaloric diets have been attributed to hormonal adaptations that override self-control of food intake. We tested this hypothesis by measuring brain fMRI reactivity to food cues and circulating energy-balance hormones in 24 overweight/obese participants before, and 1 and 3 months after starting a calorie restriction diet. Increased activity in prefrontal regions at month 1 correlated with weight loss at months 1 and 3. Weight loss was also correlated with increased plasma ghrelin and decreased leptin at month 1, and these changes were associated with greater food cue reactivity in reward-related brain regions. However, the reduction in leptin did not counteract weight loss; indeed, it was correlated with further weight loss at month 3. Activation in a network of prefrontal regions associated with self-control could contribute to individual differences in weight loss and maintenance, whereas we failed to find that the hormonal adaptations play a major role.

## Introduction

Weight loss can improve comorbidities and cardiometabolic risk factors associated with obesity. Two-thirds of the American population have undertaken reducing diets at least once (Gudzune et al., 2015). However, achieving and maintaining weight loss remain challenging (Anastasiou et al., 2015). Several studies indicate that high-order executive cognitive processes implicated in selfregulation play an important role in healthy food decisions and weight management (Gettens and Gorin, 2017; Michaud et al., 2017; Stoeckel et al., 2017). However, hormonal responses to negative energy during calorie restriction balance can modulate the activity of brain systems implicated in feeding in favor of increased calorie intake (Berthoud et al., 2012). In humans, it remains to be tested if the changes in energy balance signals during calorie restriction can modulate the brain networks associated with food intake, override self-control, and oppose weight loss.

Brain circuitry underlying food decisions can be divided into three interacting systems: (1) a homeostatic system centered around the hypothalamus; (2) a reward-related appetitive network including the striatum and the ventromedial prefrontal cortex (vmPFC) that encodes the subjective value of food cues; (3) an executive control network that relies on the function of interconnected prefrontal regions including the anterior cingulate cortex (ACC), dorsolateral prefrontal cortex (dlPFC), inferior frontal gyrus (IFG), and posterior parietal (PP) cortex (Dagher, 2012; Ochner et al., 2013). Cognitive control, defined here as the ability to restrict calorie intake and to sustain weight maintenance, is thought to rely on these executive structures. It is proposed that cognitive control ability mediates the relationship between weight loss intention and action (Gettens and Gorin, 2017). The dlPFC and IFG have been repeatedly implicated in dietary self-control, studied with functional magnetic resonance imaging (fMRI) utilizing blood-oxygen-level-dependent (BOLD) contrast. Activation of the dlPFC and IFG is seen when subjects are asked to voluntarily suppress the desire to eat in response to food cues (Batterink et al., 2010; Hollmann et al., 2012), and predicts subsequent reduced food intake outside the lab (Lopez et al., 2014, 2017). FMRI studies also support a model according to which dlPFC and IFG downregulate the activity of value-encoding regions (e.g. vmPFC) when participants choose healthy over unhealthy foods or regulate their food cravings (Hare et al., 2009, 2011). The relative balance of activity in regions associated with self-regulation over those associated with reward has been used to compute a brain-derived measure of self-regulation ability, which relates to healthier real-life food choices in dieters and non-dieters (Lopez et al., 2014, 2017). This has led to dual systems theories where behavioral outcomes depend on the balance between self-control and reactivity to reward. Although few studies have examined brain activity longitudinally in individuals undergoing calorie restriction, there is some support for the role of dlPFC in successful weight loss (Weygandt et al., 2013) and of ventral striatum activity in worse outcomes (Murdaugh et al., 2012).

According to the dual systems theory, the magnitude of weight loss during calorie restriction will be related to the following fMRI findings: (1) increased brain activity in regions associated with cognitive control, (2) increased connectivity of these areas to regions ascribed to value processing (e.g., vmPFC and striatum), and (3) downregulation of activity in these value-related brain regions. However, central nervous system networks are also modulated by internal states, such as current energy balance status (Neseliler et al., 2017; Rangel, 2013). During calorie restriction, ghrelin and leptin reflect changes in energy balance. Leptin plasma levels decline rapidly in response to calorie restriction, and more slowly with reduction of fat mass (Friedman and Mantzoros, 2015). Patients with leptin deficient states show increased food cue reactivity in the striatum and orbitofrontal cortex (OFC) compared with controls (Aotani et al., 2012). Striatal BOLD response to food cues in the striatum is reduced by leptin administration in these patients (Aotani et al., 2012; Farooqi et al., 2007). In normoleptinemic participants, leptin levels negatively correlate with food cue reactivity in the striatum (Grosshans et al., 2012) and leptin administration to weight-reduced subjects results in increased activity in regions associated with cognitive control (Rosenbaum et al., 2008). These studies suggest that reductions in leptin levels can result in increased activity in the mesolimbic reward system and reduced activity in brain regions associated with cognitive control. Conversely, ghrelin – an orexigenic hormone secreted by the stomach – increases rapidly in response to calorie deficit (Borer et al., 2009). Post-translational modification converts ghrelin to acyl-ghrelin, its active form (Müller et al., 2015). Ghrelin can increase the neural response to food cues in regions associated with value and motivation and potentiate food intake (Goldstone et al., 2014; Malik et al., 2008). During weight loss, the fall in leptin and rise in ghrelin levels could modulate the activity of brain networks involved in reward signaling to shift the balance towards increased food intake (Berthoud et al., 2012).

We designed this study to test these two predictions on the role of the central nervous system in voluntary calorie restriction: (1) that activation of cognitive control networks during fMRI would predict weight loss and (2) that early weight loss should lead to changes in energy-balance signaling (leptin and ghrelin), which would lead to increased activity in regions associated with reward and counteract self-control mechanisms. Twenty-four individuals underwent a three-month calorie restriction program with measurement of fMRI and metabolic variables at baseline (before initiating the diet), and at months 1 and 3. Body weight was also obtained at two years. During fMRI they viewed 216 pictures of appetizing foods or scenery, and rated them on a 4-point scale. Our results support the first prediction but not the second.

## Results & Discussion

### Calorie restriction resulted in weight loss

The 24 participants (1 Male) had a mean age of 37.2 (SD = ±8.4) and a mean body mass index (BMI) at entry of 30.4 (SD = ± 3.2). The personality measures of the study population are listed in Table S1. We analyzed the changes in weight, physical activity, hunger levels and energy-balance hormone levels at months 1 and 3 during calorie restriction, compared with baseline (Fig. 2). All the analyses were conducted using linear mixed effect modeling. Significant reductions in BMI occurred across the three sessions of calorie restriction (F (1,62) = 86.2, p = 2.4 *10^−13^). Pairwise comparisons showed the reductions in BMI were significant from baseline to month 1 (F (1,61) = 22.91, p = 1.12 *10^−5^, mean weight loss 2.20 ± 4.22 kg), and from month 1 to month 3 (F (1,61) = 17.16, p = 0.0001, mean weight loss 1.88 ± 1.55 kg) (Fig. 2-A, Table S2). Self-reported physical activity levels across the sessions did not show significant differences (F (1,31) = 0.31, p = 0.71, Table S2). There were no significant differences in hunger assessed by VAS across the sessions (Table S3).

**Figure 1.**
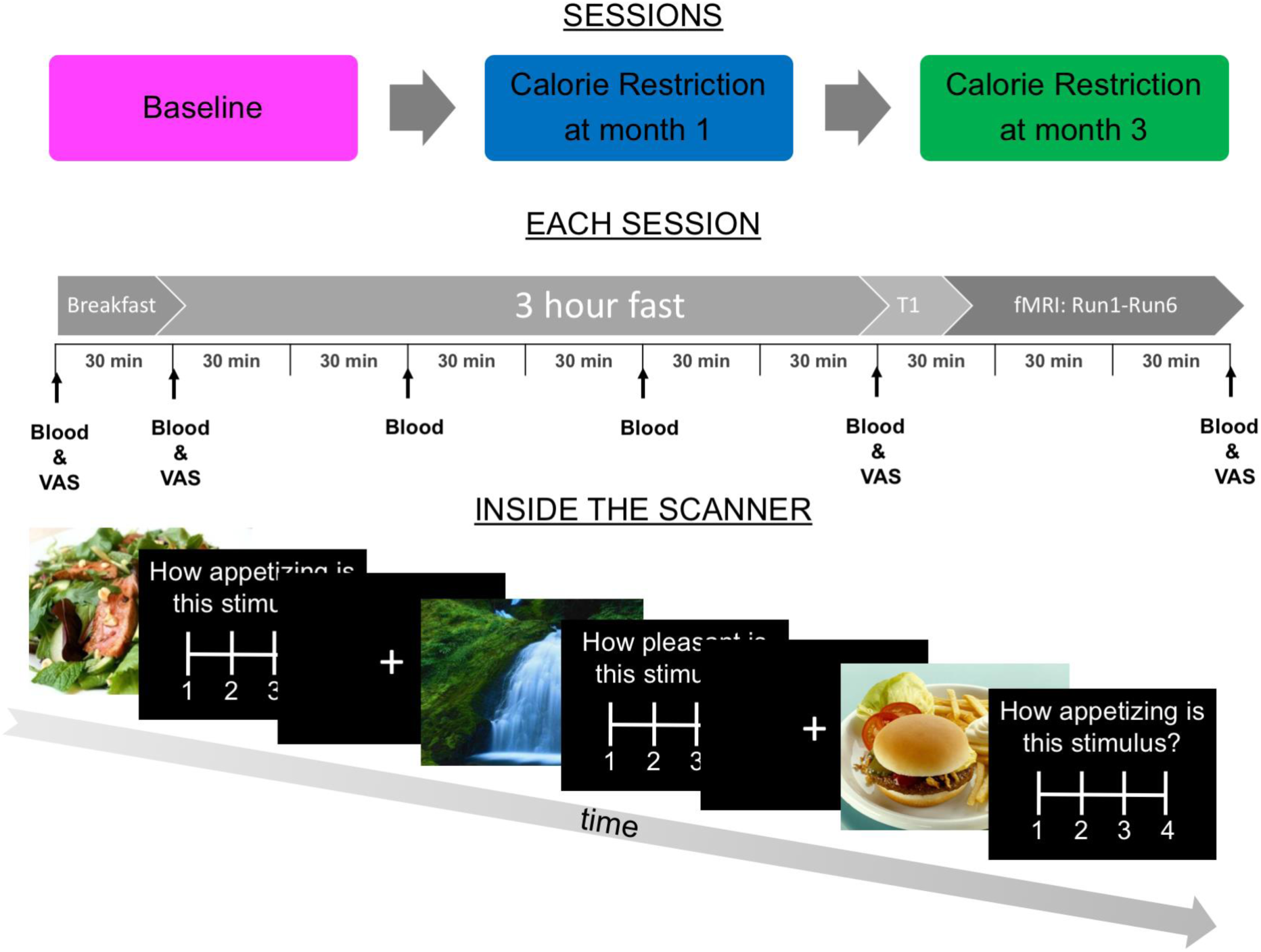
Experimental Design. Participants (n=24) were tested three times: at baseline, then at 1 and 3 months during voluntary calorie restriction. On each scan day, participants ate a standard breakfast three hours prior to fMRI imaging. Venous blood and self-reported hunger level on a visual analog scale (VAS) were sampled at the times indicated. Anatomical scanning (T1) was followed by six seven-minute functional runs. During fMRI participants viewed pictures of foods and scenery followed by a rating.

**Figure 2.**
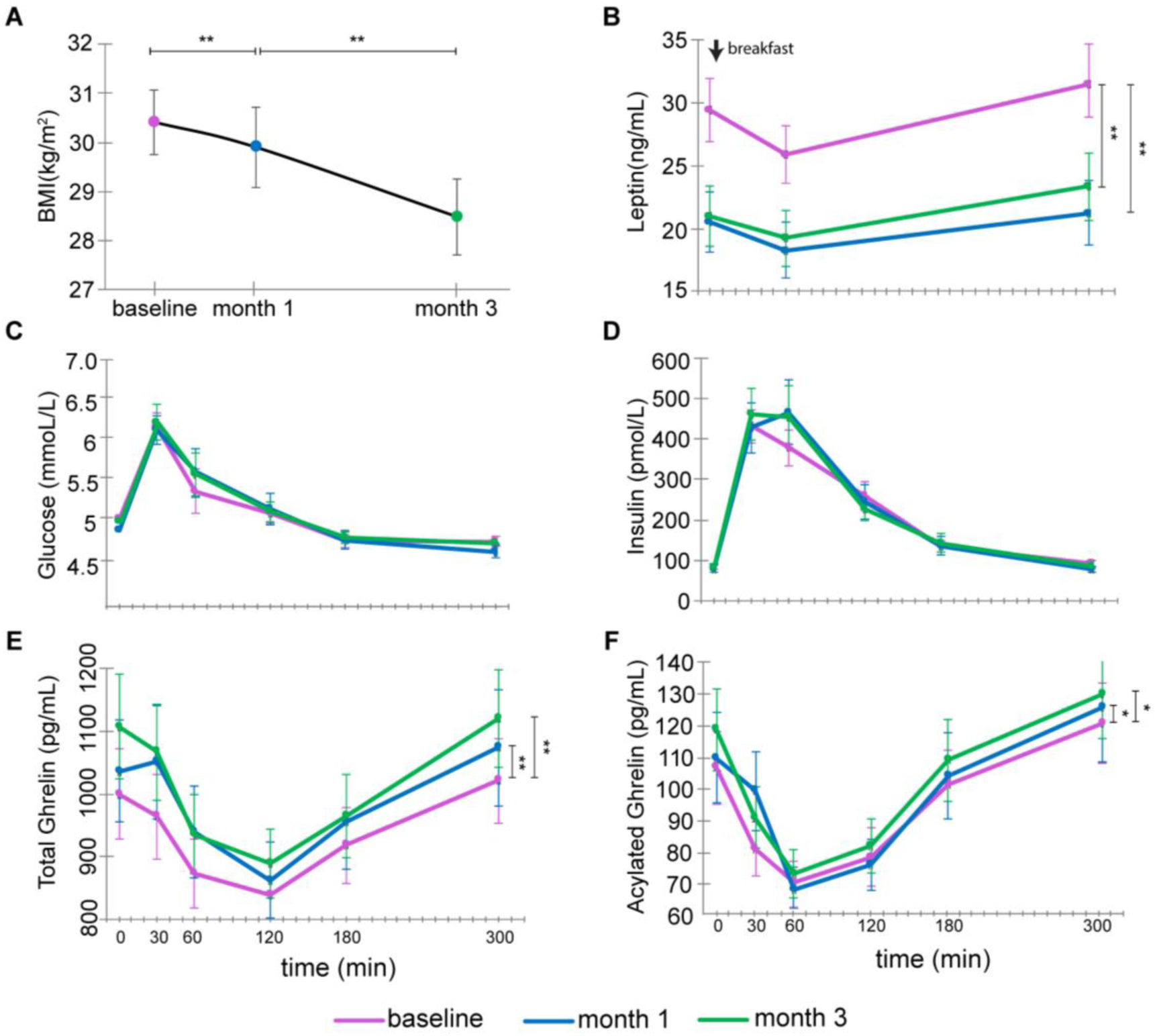
Effects of voluntary calorie restriction. (A) BMI decreased significantly across the sessions. In panels B-F, the plasma concentrations were measured throughout the experiment (0-300 min). 0 min refers to morning (pre-meal) levels, after which participants consumed the standard breakfast (shown as an arrow in B). In all the panes, 30 min indicates the response after the breakfast. (B) Leptin decreased at month 1 during the diet compared to baseline, but was not significantly different at month 3 compared with month 1. Glucose (C) and Insulin (D) did not show significant differences across sessions. Total ghrelin (E) and acylated ghrelin (F) levels were lower at month 1 and month 3 compared with baseline. Data are presented as mean ± SEM. Statistics are derived from linear mixed models (MATLAB function fitlme). * = p <0.05; ** = p<0.01.

### Activity in cognitive control networks correlated with weight loss

We hypothesized that weight loss would be related to activity in regions implicated in cognitive control. In line with this hypothesis, initial weight loss at month 1 was correlated with an increase in BOLD (month 1 versus baseline) during the food minus scenery contrast in regions associated with cognitive control such as dlPFC, IFG, dACC, inferior parietal lobule, and caudate (Fig. 3-A&B, Table S5). This result is in line with studies that showed that cue-related dlPFC activity correlated with weight loss from dieting (Weygandt et al., 2013) or bariatric surgery (Goldman et al., 2013). Food cue-reactivity in the network of regions related to cognitive control at month 1 (Fig 3-A) correlated positively with subsequent weight loss from month 1 to 3 (r = 0.60, p = 0.013, Fig. 3-C). Moreover, activity reductions in this network at month 3 (i.e. a return towards baseline) correlated with weight regain two years later (r = −0.64, p=0.014, Fig. 3-D). These results suggest that engagement of prefrontal areas implicated in dietary self-control is correlated with initial weight loss at months 1 and 3 and weight loss maintenance at two years.

**Figure 3.**
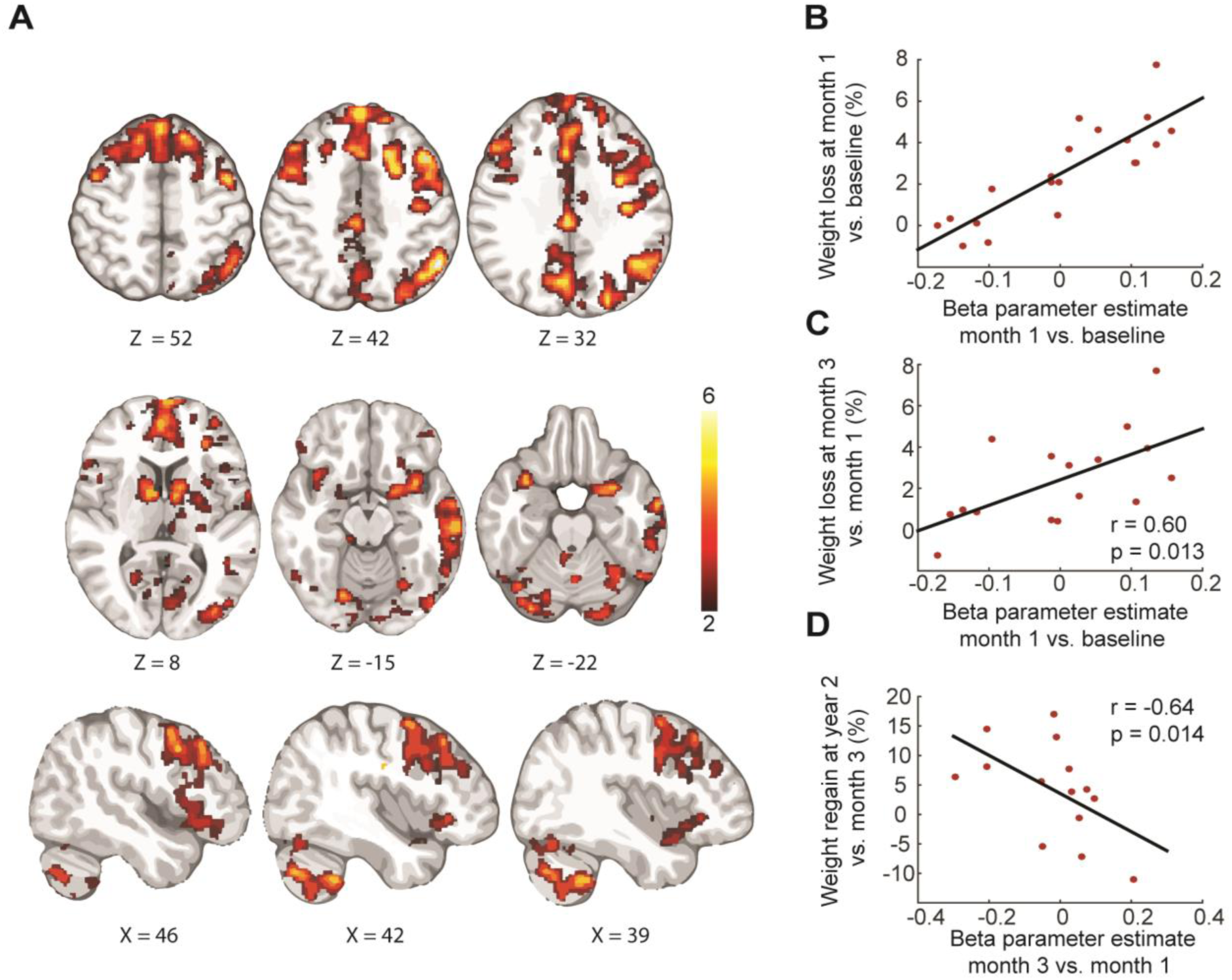
Weight loss at month 1 correlated with changes in BOLD in regions associated with cognitive control. (A) Activation to food cues compared to scenery cues at month 1 vs. baseline correlated with weight loss (p<0.05 FWER, N=20). (B) Mean beta estimate of activation to food minus scenery at month 1 minus baseline derived from the significant cluster (A) versus weight loss between month 1 and baseline. (C) Mean beta estimate of activation to food minus scenery at month 1 minus baseline derived from the significant cluster (A) versus weight loss from month 3 to month 1 (N = 16). (D) Mean beta estimate of activation to food minus scenery at month 3 minus month 1 derived from the significant cluster (A) versus weight regain at two years compared with month 3 (N = 14). The color scale presents t-statistics derived from 5000 permutations of the data. FWER: Family-Wise Error Rate. X and Z refer to the MNI coordinates in mm.

### vmPFC BOLD decreased during calorie restriction

Neuro-computational models of decision-making assume that individuals make choices based on the subjective value assigned to options (Rangel, 2013). As an example, a subjective value signal for food stimuli might be computed by incorporating aspects of palatability and healthiness. Meta-analysis reveals that this stimulus value computation is reflected in the vmPFC (Bartra et al., 2013), and influenced by its connectivity with regions implicated in reward processing, such as the striatum, and cognitive control, such as the dlPFC and IFG (Hare et al., 2009, 2011; Rangel, 2013). We tested two competing predictions regarding the change in vmPFC response to food cues during calorie restriction: (1) it could increase, reflecting greater valuation of food cues due to negative energy balance; (2) it could show a reduction, reflecting successful self-regulation (Hare et al., 2009). Food cue reactivity was reduced in vmPFC at month 1 compared to baseline in a region of interest (ROI) derived from a meta-analysis of subjective value (Bartra et al., 2013) (Fig. 4-A; Table S6A). Activity in vmPFC in the food minus scenery contrast remained lower than baseline at month 3 (Fig. 4-B) in line with continued weight loss. There was no difference in vmPFC activation (food minus scenery) at month 3 compared to month 1.

**Figure 4.**
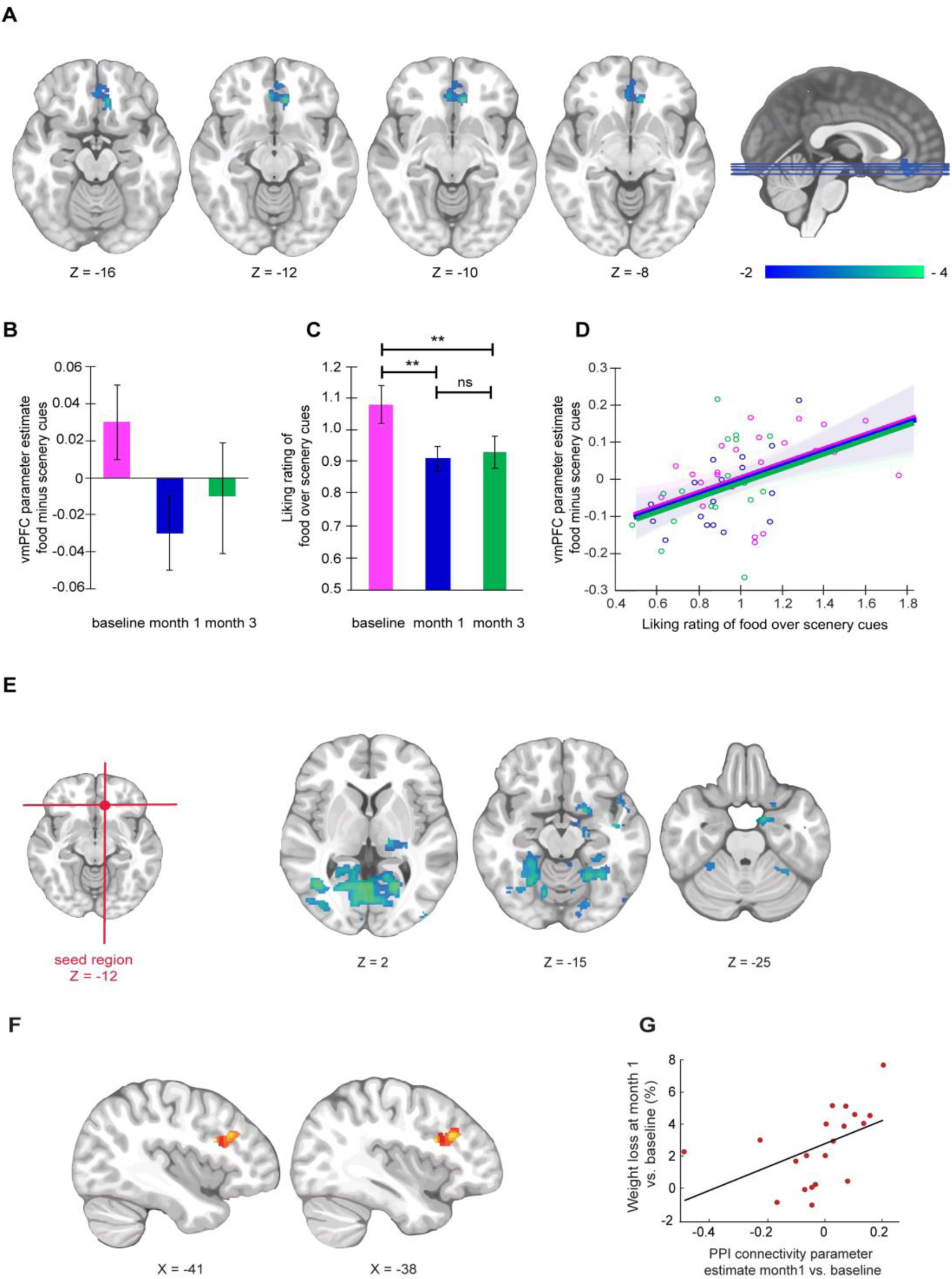
vmPFC activity following calorie restriction. (A) Negative peaks: activation to food cues compared to scenery cues at month 1 of the diet compared to baseline (p<0.05 FWER, SVC, N = 20). (B) Changes in the vmPFC region beta estimate derived from the entire cluster in A. (C) Liking for food cues relative to scenery cues at baseline, month 1 and month 3. (D) Mean vmPFC parameter estimate derived from the entire cluster to food minus scenery cues versus mean liking ratings of food cues relative to scenery cues (F(1,61) = 13.24, p = 0.0006). Shaded lines represent 95% confidence intervals derived from the linear mixed effect model (MATLAB fitlme). (E) PPI analysis with the vmPFC seed revealed that left vmPFC connectivity is reduced with visual areas at month 1 compared with baseline (displayed p< 0.001, uncorrected, minimum voxel extent = 10 mm, p<0.05 FWER corrected in Table S6-C). (F) Left vmPFC connectivity to left dlPFC and left IFG at month 1 compared with baseline is positively correlated with weight loss (displayed p< 0.001, uncorrected, minimum voxel extent = 10 mm, p <0.05 FWER corrected in Table S6-D). (G) The mean beta estimate derived from the significant cluster (F) in month 1 compared with baseline versus weight loss between month 1 and baseline. Data are presented as mean ± SEM and statistics are derived from linear mixed models. SVC: Small Volume Correction. FWER: Family-wise Error Rate. ns: not significant. ** = p<0.01. X and Z refer to MNI coordinates in mm.

### Food liking decreased during calorie restriction

Subjective liking ratings for food cues relative to the scenery cues showed a similar trend: there was a significant reduction at month 1 (F(1,61) = 283.33, p = 1.34 * 10^−24^) that persisted at month 3 compared with baseline (F(1,61) = 273.73, p = 0.0012). Month 3 liking ratings were not significantly different from month 1 (F(1,61) = 0.23, p= 0.63) (Fig. 4-C). These result align with previous observations showing that food cravings subside during voluntary calorie restriction (Martin et al., 2006). Furthermore, liking for food vs. scenery cues correlated with vmPFC food cue reactivity (linear mixed effects model F(1,61) = 13.24, p = 0.0006) across all sessions (Fig. 4-D), supporting a role for this region in value computation. In sum, both fMRI vmPFC signals and liking for food cues were reduced during calorie restriction, consistent with previous studies linking these to self-regulation of appetite (Hare et al., 2009; Wagner et al., 2013).

### vmPFC connectivity changed during calorie restriction

The connectivity of the vmPFC (seed region MNI coordinates: x = −10, y = 34, z = −12, Fig. 4-E), was reduced at month 1 vs baseline with regions associated with visual processing of food stimuli. These regions included the lingual gyrus, the lateral and temporal occipital cortex regions (Fig. 4-E, Table S6B). Focusing on the visual features of valued stimuli has been associated with BOLD response in these regions (van der Laan et al., 2011; Lim et al., 2011), which have been postulated to send the visual information to the vmPFC, where an overall subjective value of the stimulus is computed and utilized for decision making (Lim et al., 2013). Weight loss also correlated with increased vmPFC connectivity to left dlPFC in regions previously associated with cognitive control (by using a mask generated from the search term “cognitive-control” in the meta-analytical tool Neurosynth) (Fig.4-F, Table S6C).

### IFG activity subsided from month 1 to 3 during calorie restriction

These effects tended to subside from month 1 to 3, when there was a reduction in activity in right IFG/Frontal Opercular Cortex (Fig. S1-A; Table S6D), and reduced vmPFC connectivity to left dlPFC (Fig. S1-B; Table S7). Participants who showed more reduction in right IFG BOLD at month 3 vs. month 1 showed greater weight-regain at 2-year follow-up (r = −0.61, p =0.02, n=14, Fig. S1-B).

These results support a model in which calorie restriction entails changes in value computations in vmPFC due to reduced connectivity with visual areas involved in computing attributes of stimuli (Lim et al., 2013) and increased connectivity with prefrontal areas implicated in self-control (Rangel, 2013). These results build on previous work showing that dlPFC activity and its connectivity to vmPFC correlates with self-regulation and reduced food consumption (Hare et al., 2009; Lopez et al., 2014). In this study, changes in brain activity in these regions predicted subsequent real-life outcomes and correlated with the magnitude of weight loss. However, we observed that both the activity in regions associated with cognitive control as well as their connectivity to vmPFC were reduced at month 3 compared with month 1 suggesting that BOLD changes in cognitive control networks may not be sustained, which appears to correlate with subsequent weight regain (Fig. 3-D).

### Leptin decreased, and ghrelin increased during calorie restriction

Leptin and ghrelin both showed adaptations to weight loss. Plasma leptin showed significant effects for both session (F (1,188) = 81.91, p = 1.79 * 10^−16^) and sampling time (F (1, 188) = 7.29, p = 0.008) (Fig. 2-B). Levels decreased from baseline at both month 1 (F (1, 188) = 100.52, p = 3.27 * 10^−19^) and month 3 (F (1, 188) = 88.27, p = 2.03 * 10^−17^), with no change between month 1 and 3 (F (1, 188) = 0.34, p = 0.56). There was a decrease from time 0 to 60 min after breakfast (F (1, 187) = 8.10, p = 0.005), followed by an increase post-scan (F (1, 187) = 19.04, p = 2.12 * 10^−5^), to levels not significantly different from baseline (F (1, 187) = 2.30, p = 0.13).

Total plasma ghrelin levels showed a main session effect with increase at month 1 (F (1,379) = 27.58, p = 2.52 *10^−7^), remaining high at month 3 (F(1,379) = 26.48, p = 4.29 *10^−7^), with no change between months 1 and 3 (F (1,379) = 0.017, p = 0.90) (Fig. 2-E). Levels decreased to a nadir at 120 min after the breakfast (F (1,376) = 46.5, p = 4.54 * 10^−11^) then rising, the increase being significant from 120 to 180 min (F (1,376) = 18.01, p = 2.77 * 10^−5^), yet lower than at time 0 (F(1,376) = 46.5, p = 4.54 * 10^−11^). During the scan ghrelin increased further (F (1,376) = 41.81, p = 3.11 * 10^−10^), to a level not different from time 0 F (1,376) = 1.6, p = 0.21).

Plasma acyl-ghrelin (Fig.2-F) showed a main effect of increase at month 1 (F (1,379) = 5.00, p = 0.026), remaining elevated at month 3 relative to baseline F (1,379) = 4.27, p = 0.039). They were not different between months 1 and 3 (F (1,379) = 0.025, p = 0.87). As expected, levels decreased markedly by 60 minutes following breakfast (F (1,376) = 22.63, p = 2.81* 10^−6^) then rose progressively thereafter. Levels at the end of scan (300 min) were not different than at time 0 (F (1, 376) = 2.2, p =0.14), but increased during the scan (180 vs. 300 min) (F (1, 376) = 20.39, p = 8.46 * 10^−6^).

The magnitude of month 1 weight loss was correlated with pre-meal log leptin and ghrelin levels. It was negatively correlated with pre-meal log leptin (r= − 0.78, p = 1.56 * 10^−5^, Fig. 5-A), and positively with pre-meal total ghrelin levels at month 1 (r = 0.67, p = 7.41 * 10^−4^, Fig. 5-B) but not with acyl-ghrelin levels (r = 0.20, p = 0.38). The effects remained at month 3 when weight loss compared with baseline correlated inversely with pre-meal leptin levels (r = −0.86, p = 5.39 *10^−6^) and positively with pre-meal acyl ghrelin (r = 0.47, p = 0.048) and total ghrelin levels (r = 0. 56, p = 0.016).

**Figure.5.**
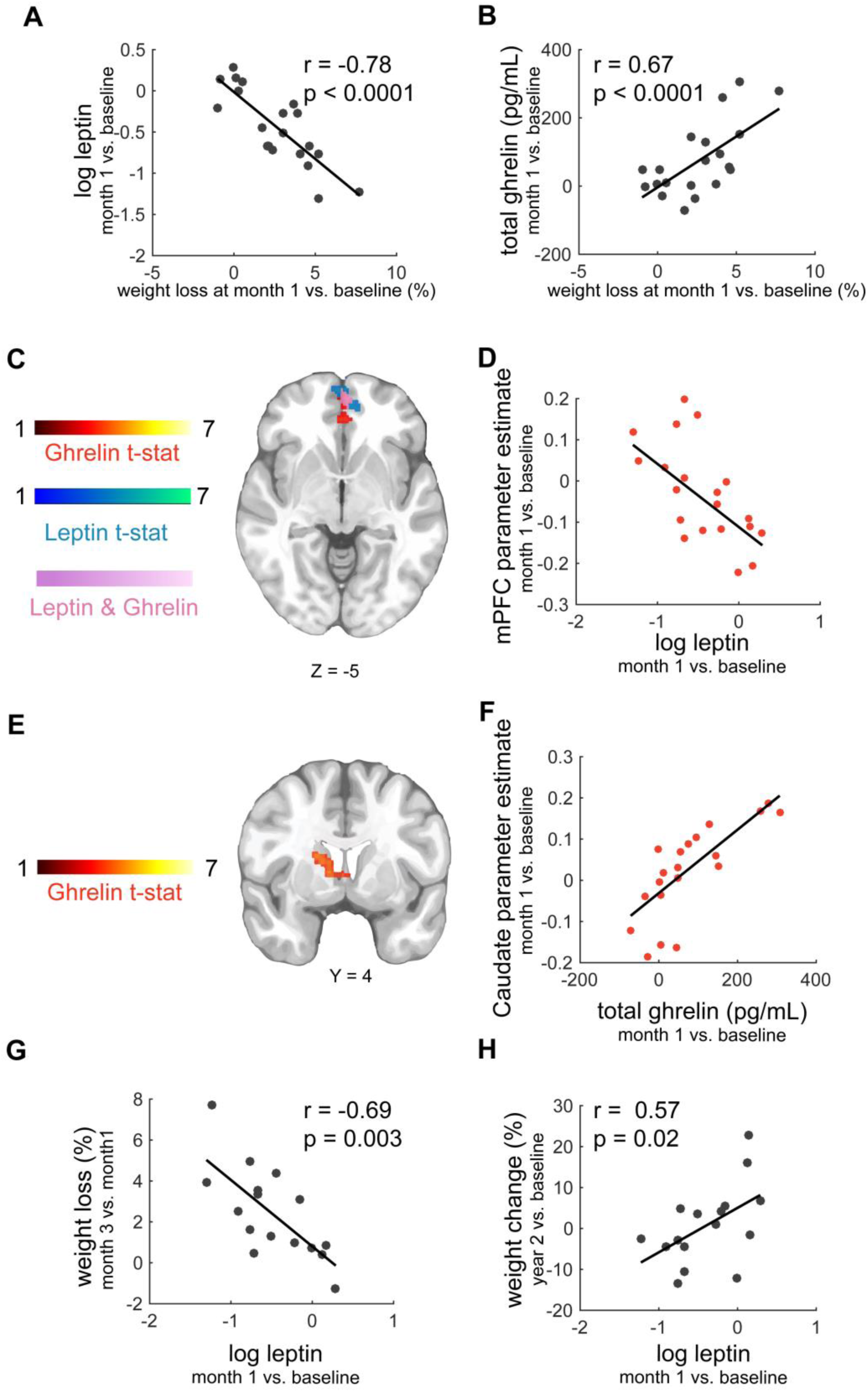
Correlations with leptin and ghrelin levels. (A) Weight loss at month 1 vs. baseline versus pre-meal plasma leptin and (B) total ghrelin at month 1 vs. baseline. (C) Correlation of food minus scenery activation at month 1 vs. baseline and the increase in pre-meal ghrelin levels (orange); and the reduction in pre-meal log leptin levels (blue) at month 1 vs. baseline. Pink regions display areas significant in both analyses (displayed p <0.05 FWER, Table S6A). (D) Depiction of the correlation from panel C. The mean beta parameter estimate is derived from the orange cluster in panel C at month 1 vs. baseline. (E) Correlation of food minus scenery activation at month 1 vs. baseline and the increase in pre-meal ghrelin levels (orange); (displayed p <0.05 FWER, Table S6A). (F) Depiction of the correlation from panel E. The mean beta parameter estimate was derived from the orange cluster in panel E (G) Pre-meal log leptin levels at month 1 vs. baseline correlated inversely with future weight loss at month 3 vs. month 1 and (H) negatively with weight at two years compared with baseline. The color scales represent t-statistics derived from 5000 permutations of the data. FWER: Family-Wise Error Rate. Y and Z refer to MNI coordinates in mm.

No significant effects of session on glucose and insulin levels were detected (Fig. 2 C-D). In addition, insulin sensitivity as assessed by homeostatic model assessment (HOMA) was not significantly different across sessions (F(1,61) =1.37, p=0.25, Table S2). These results are consistent with insulin resistance playing a minor role in the main outcome variables in this study.

Leptin is produced by adipocytes. Its plasma levels fall quickly during negative energy balance, and more gradually with diminished fat mass (Friedman & Mantzoros, 2015). The orexigenic hormone ghrelin is increased in response to energy deficit (Borer et al., 2009; Müller et al., 2015). In the current study, leptin levels decreased while ghrelin levels increased at month 1, consistent with previous weight loss studies (Wing et al., 1996). Despite continued weight loss, ghrelin and leptin levels did not change further at month 3, as described previously (Crujeiras et al., 2010; Sumithran et al., 2011) These results suggest that, on average, our participants were in negative energy balance at month 1 and month 3 compared with baseline.

### Ghrelin and leptin changes correlated with BOLD in reward regions

We tested the hypothesis that weight loss-induced changes in ghrelin and leptin would increase brain activation of reward related areas while viewing food cues, and that this would promote overeating and diet failure. Previous studies found that leptin administration reduced food cue reactivity in leptin deficient patients (Aotani et al., 2012; Farooqi et al., 2007); while ghrelin administration increased food cue reactivity (Malik et al., 2008) in regions associated with food value and motivation.

Consistent with previous studies (Goldstone et al., 2014; Kroemer et al., 2013; Malik et al., 2008), increased pre-meal ghrelin levels correlated with increased activity at month 1 in mPFC, caudate and visual cortex, all areas associated with appetitive processing (Fig. 5C,E-F; Table S8). Weight loss-induced reductions in pre-meal leptin level also correlated with increased activity in mPFC and visual cortex (Fig. 5C-D, Table S9), suggesting that metabolic adaptations at month 1 of contributed to increased food cue reactivity at month 1 compared with baseline.

### Ghrelin and leptin changes did not oppose continued weight loss

The next question we addressed was whether negative energy balance signals could counteract participants’ efforts to continue losing weight by increasing value-related food cue reactivity and food intake. Contrary to our hypothesis, leptin reductions at month 1 vs. baseline correlated with further weight loss at month 3 vs. month 1 (r = −0.69, p = 0032, Fig. 5-G) and a lower weight at two years compared with baseline (r = 0.57, p = 0.02, Fig. 5-H). Increases in ghrelin levels did not significantly relate to weight change at 3 month (r = 0.43, p = 0.099), or at two years compared with baseline (r= − 0.34, p = 0. 20), however the direction of the effect was not consistent with ghrelin causing an increase in weight. These results suggest that leptin reductions and ghrelin increases did not oppose continued weight loss, even though they increased reward related food cue-reactivity. Indeed, leptin reductions predicted greater subsequent weight loss. This is likely because leptin reductions were greater in individuals with better appetite control, who lost more weight.

In this study, leptin and ghrelin adaptations to weight loss did not explain why diets are unsustainable in the long term. Our results suggest that the reduction of leptin at month 1 are a marker of successful weight loss, correlating with future weight loss and/or maintenance. Indeed, in the Look Ahead study, weight loss at month 1 predicted reduced weight up to 8 years later (Unick et al., 2015). Our results are not consistent with the prevalent view that reduced leptin levels during energy restriction promote increased food intake and obesity, a theory that has been recently questioned (Flier and Maratos-Flier, 2017). A recent meta-analysis of reduced leptin levels as a predictor for weight regain was also inconclusive (Strohacker et al., 2014) and leptin pharmacotherapy in the treatment of obesity has produced disappointing results (Bray, 2014; Korner et al., 2013). All together, these results do not support pharmacologically increasing leptin signaling as an adjunct target for treatment of obesity in conjunction with calorie restriction. However, more research is needed to test the role of leptin for the treatment of obesity in normoleptinemic patients (Friedman and Mantzoros, 2015; Rosenbaum and Leibel, 2014).

## Conclusions

Weight loss is highly variable among individuals embarking on weight-reduction programs and likely depends on interactions between peripheral and central mechanisms of appetite control. While changes in short- and long-term energy signals due to weight loss can affect food intake to maintain a stable weight (Ryan, Woods, & Seeley, 2012), in humans food intake is also under the influence of cognitive goals and self-regulation. It is important to note that cognitive control changes were not measured here but inferred from the activation of lateral prefrontal areas while viewing food items. Nonetheless, our principal findings are that changes in neural correlates of cognitive control from baseline to month 1 were associated with the magnitude of weight loss at months 1 and 3, while reduced activity from months 1 to 3 in these regions correlated with subsequent weight regain. In addition, increased ghrelin and reduced leptin levels were observed during calorie restriction, but these responses, while increasing food-cue reactivity in reward related brain regions, did not counteract further weight-loss. Thus, we failed to show that hormonal adaptations to negative energy balance play a significant negative role in weight loss success and its maintenance. Therefore, it is possible that strategies that target the central brain networks involved in cognitive control might be more effective as weight loss strategies.

## Limitations

Although our small sample size is common for within-design neuroimaging studies, it limits our statistical power and the generalizability of our findings. In addition, for some participants we utilized self-reported weight at 2 years, which might have biased our results. Second, we only tested total ghrelin, acyl-ghrelin, leptin, glucose and insulin levels. Therefore, we cannot generalize our results to other metabolic signaling molecules such as glucagon-like peptide (GLP-1). Third, the communication between the metabolic hormones and the brain might have changed as a function of weight status (Ravussin et al, 2014); e.g., leptin resistance might have been present in our sample. We did not test for hormonal resistance, and therefore cannot determine how much changes in peripheral signaling affected brain responses at different time points. Nonetheless, leptin and ghrelin did have the expected effects on cue-reactivity in appetitive brain regions. Another limitation is the lack of a control group. Further studies are justified, and could include larger cohorts, a control group, monitoring of daily food intake and activity, indirect calorimetry, more precise estimation of insulin sensitivity, and additional known modulators of food intake and activity.

## Acknowledgments

We thank Dr. Isabelle Garcia-Garcia and Dr. Andreanne Michaud for critically reading the manuscript. AD is supported by Canadian Institutes of Health Research (CIHR) Foundation Scheme. SN is supported by the Frederick Banting and Charles Best Canada Graduate Scholarship (CIHR). This projected was funded by CIHR Operating Grant number 219271.

## Author Contributions

Conceptualization, S.N, W.H., E.B.M and A.D.; Methodology, S.N., W.H., K.L, M.L., E.B.M., M.D. and Y.Z.; Software, S.N., M.D. and K.L.; Investigation, S.N., W H, M.Z., S.G.S., M.L., S.C.S, M.L.; Writing – Original Draft, S.N. and A.D.; Writing – Review & Editing, S.N., M.D., E.B.M. and A.D.; Funding Acquisition, E.M. and A.D.; Resources, E.B.M., M.L., S.S. and A.D.; Supervision, E.B.M. and A.D.

## Materials and Methods

### Participants

29 right-handed participants [1 male, BMI = 30.94 (SD = ±3.76); age = 37.28 (SD = ±7.99)] were recruited in a private weight loss clinic (“Clinique Motivation Minceur”, Montréal. QC Canada) before they started a prescribed weight-loss regimen. After the protocol was explained, those accepting to participate signed the consent form, underwent medical history, physical examinations and blood testing to look for comorbid conditions. Inclusion criteria were: healthy apart from obesity, and stable weight at least for three months. Exclusion criteria were diabetes, uncontrolled hypertension, currently smoking, substance abuse and current use of a central nervous system active medication, renal disease, non-dermatologic cancer in the previous 5 years and current or history of neurological, eating or psychiatric disorders. Candidates who could not undergo MRI due to claustrophobia, pregnancy, implanted metal, or BMI>40 kg/m^2^ were also excluded (n = 1). Each participant received an individualized non-ketogenic calorie-restricted diet program from the weight loss clinic. The prescribed diets contained 1100-1400 kcal/day, with 40% carbohydrate, 30% fat and 30% protein. Oral calcium and multivitamins were recommended, as was 30-45 min of brisk walking per day. Weekly visits included motivational counselling. The study was approved by the Montreal. Neurological Institute Research Ethics Board. Volunteers received compensation for participation in the form of free nutritional supplements and reimbursement for travel.

Data from one participant was not included due the presence of uncorrectable fMRI brain artifacts (n = 1). Lack of button response to more than 25% of the stimulus ratings deemed a run incomplete as it indicated that the participant was not paying attention to the task. Any session with 50 % incomplete runs (3 out of 6 runs) was excluded. Three participants were inattentive during the fMRI scan during at all 3 sessions (n = 3). In addition, four did not successfully complete the fMRI at session 2 (leaving n=20) and four (others) at session 3 (leaving n = 20). This reduced the sample size for comparing baseline to month 1 to 20 participants, and the comparison of month 1 to month 3 to 16 participants. Participants were contacted at two years following the experiment. Of 19 respondents, weights were reported by 10, and measured in 9. Of these 19 respondents, 14 people completed all three sessions of the fMRI successfully. Weight at two years is calculated as the percent change of weight from baseline to two years (n = 19); and weight regain as the percent change of weight from month 3 to two years (n = 14).

### Experimental Design

Participants underwent fMRI on three occasions: first immediately prior to initiation of the diet program, then at month 1 and month 3 during calorie restriction. Women were scanned during the luteal phase of their menstrual cycle. The scanning sessions started between 12:00 and 1:30 PM and kept the same schedule for the three sessions. On the scan days, participants presented themselves to the lab in the morning, having fasted from midnight the night before. All participants received the same standardized breakfast. This included ¾ cup of 2% milk, 1 slice of brown toast, 10 mL of peanut butter, one medium-sized hard-boiled egg, one small apple, a protein bar and 150 mL of black coffee or tea without sugar. It contained 550 kcal, 43% carbohydrate, 32% fat and 25% protein and was eaten over 10-15 minutes. Venous blood samples were drawn through a stopcock in an antecubital vein immediately prior to breakfast, then at 30, 60, 120 and 180 min (just prior to the fMRI scan) then just after the scan at 300 min (Fig.1) to measure hormone levels (see below). In addition, at each session, blood tests were conducted to measure C reactive protein, ketones, cholesterol, triglyceride, and glycated hemoglobin (HbA1c) levels (Table S2). Participants were confirmed to have negative blood ketones at each session.

### Psychological, Physical Activity, Hunger Measures

Participants completed the Binge Eating Scale, the Dutch Eating Behavior (van Strien et al., 1986) and Power of Food Questionnaires (Lowe et al., 2009) to assess the eating styles. Sensitivity to reward and punishment was assessed using the Sensitivity to Punishment and Sensitivity to Reward Questionnaire(Torrubia et al., 2001) and the BIS/BAS Scale (Carver and White, 1994), depression levels using the Beck Depression Inventory(Beck A.T., 1972)), and chronic stress during the previous month using the Perceived Stress Scale (PSS) (Cohen et al., 1983) (Table S1). Participants reported their activity levels over the past week by answering two questions: “Using the past week a reference, how much time have you given to the following activities? Jogging, cycling, swimming, cross-country skiing, aerobic dance or other similar activities.” and “Over the past week, how much time have you spent doing strenuous work, a physical activity or a sport other than those mentioned in the previous question?” The mean scores from these two questions were used to calculate changes in self-reported exercise (Table S2). Hunger levels were assessed using Visual Analog Scales (VAS) four times throughout the day across the sessions (Fig. 1). On the VAS, we asked the participants “On a scale from 0 to 10 how hungry do you feel now?” (Table S3).

### Hormone Measurements

The blood samples were centrifuged, and plasma and serum stored at −80C until analysis. All samples from each participant were measured within the same assay. Plasma glucose was measured by the glucose-oxidase technique (GM-9, Analox Instruments, Lunenberg, MA, USA), and insulin using a specific radioimmunoassay (RIA) (Millipore, Billerica, MA, USA). We measured both total and acylated ghrelin, as the latter is considered “active”. Samples required acidification before storage to prevent degradation (Hosoda and Kangawa, 2012). Both were measured by RIA (Millipore, Billerica, MA, USA). Leptin was assayed using a solid phase ELISA (R&D Systems Inc, Minneapolis, MN, USA). The HOMA-IR (insulin resistance) index was calculated as fasting (glucose x insulin)/22.5.

### MRI

Neuroimaging was carried out with a Siemens Magnetom Trio 3T MRI scanner at the Montreal. Neurological Institute (MNI). High-resolution T1-weighted anatomical images with voxel size = 1x1x1 mm were obtained first. Functional data were acquired with an echo-planar T2*weighted sequence for BOLD contrast (TR = 2 s; TE = 30 ms; flip angle, 90°; FOV = 224mm, voxel size = 3.5 × 3.5 × 3.5 mm^3^, number of slices = 38).

For each session, participants underwent six 7-minute functional runs. During each run subjects viewed images of food or scenery, presented via a projector and a mirror placed on the head coil. The food and scenery images (examples in Fig.1) had been previously matched for visual appeal (Malik et al., 2008). Each run comprised 36 unique images, 12 each of high and low-calorie foods (e.g. brownies, vegetables) and scenery (Fig.1). The order of picture presentation was randomized across subjects and runs. Images were presented for 4.0 seconds and were followed by a rating of the stimulus on a 1-4 scale. For food pictures, participants rated “How appetizing is this stimulus?” and for scenery pictures participants rated “How pleasant is this stimulus?”. Rating was followed by a fixation cross with a jittered interstimulus interval (2.5-6.0 seconds). Stimulus presentation was done using E-Prime (Psychology Software Tools Inc, Sharpsburg PA, USA). Ratings were entered by subjects via a MR-compatible button device.

## Quantification and Statistical Analysis

### Hormone and Behavioral Analysis

The values and descriptive statistics for the hormone measurements are listed in Table S3. The statistical analysis of hormonal and psychological measures was conducted using MATLAB (Version R2015a, The MathWorks Inc., Natick, MA, USA). We log-transformed leptin and insulin values to correct for non-normality. The longitudinal analyses were run using linear mixed effects modelling (MATLAB function fitlme) with subject as a random effect. The Akaike information criterion was utilized to select the best model (Table S4). To compare across sessions and across time points, and to calculate the F and p values, we conducted linear hypothesis testing on the linear mixed regression model coefficients using CoefTest implemented in MATLAB. The graphs were produced using the raw data, but the statistical analyses reported were based on linear mixed effect models. For correlations, we conducted Spearman rank correlations when the distribution was not normal.

### Imaging Data Analysis

The T1-weighted MRIs were submitted to brain extraction using BEaST, a nonlocal segmentation method applied to the images linearly registered to ICBM-MNI template (Eskildsen et al., 2012). Pre-processing of the BOLD data was conducted using FEAT (FMRI Expert Analysis Tool, Version 6, part of FMRIB’s Software, www.fmrib.ox.ac.uk/fsl) and consisted of slice timing motion correction, spatial smoothing (6mm), and high-pass filtering with a cut-off frequency of 0.1 Hz (Jenkinson et al., 2012). Linear registration (6 parameters) to T1-weighted and non-linear registration to standard T1-weighted ICBM-MNI152 template brain (voxel size= 2×2×2 mm^3^) were completed using FLIRT (Jenkinson et al., 2002) and FNIRT (Andersson et al., 2007) respectively, prior to statistical analysis.

### Imaging Statistics

For the first-level statistical analysis of the BOLD time-series, a general linear model (GLM) was implemented using FILM (Woolrich et al., 2004).The regressors for the GLM were the motion parameters, button presses for ratings, missed events, and the food and scenery image presentation events which were modeled by convolving the time course with a double-gamma hemodynamic response function (HRF) and applying temporal filtering. The resulting contrast images (i.e. food minus scenery) were then passed onto a second-level fixed-effects analysis for each subject, to obtain mean contrast estimates over runs within subjects by forcing random effects variance to zero in FLAME (FMRIB’s Local Analysis of Mixed Effects) (Beckmann et al., 2003; Woolrich et al., 2004) The t-stat maps for the contrast of interest (i.e., session1 vs. session2, session2 vs. session3, etc.) were combined across subjects for the third-level analysis.

In the third level analysis, the results at the group level were analyzed using non-parametric permutation tests (n = 5,000 permutations) with the threshold-free cluster enhancement (TFCE) algorithm using randomize in FSL (Smith and Nichols, 2009). In order to explore how the percent weight loss or changes in hormone levels affected the reactivity to food cues (i.e. food vs. scenery images), we included them as regressors in the GLM model. We utilized percent change in weight loss to account for baseline weight.

To analyze the effects of weight loss on value related activity for food items, we performed a region of interest analysis using a mask from a fMRI meta-analysis for subjective value at the time of decision (see Fig.6A from (Bartra et al., 2013)). In addition, we investigated how weight loss related to changes in BOLD in regions implicated in cognitive control. For this purpose, we restricted our analysis to a regional forward-inference mask associated with the term “cognitive control” in the NeuroSynth database of fMRI studies (http://neurosynth.org/analyses/terms/cognitive%20control/) (Yarkoni et al., 2011). Featquery from FSL was utilized to extract the percent change in mean beta parameter estimates in each ROI that was used for plotting.

### Functional Connectivity Analysis

For functional connectivity analysis we utilized psychophysiological interaction (PPI) (O’Reilly et al., 2012). We selected a seed region based on the peak coordinate of food minus scenery at session 1 minus session 2 (MNI: x = −10, y = 34, z = −12). This peak was used to form a 4 mm diameter seed region of interest (ROI) in the vmPFC. We further transformed this ROI in standard space into each individual’s functional space using the previously obtained linear transformations (from FLIRT). For each subject, we determined the peak voxel within the ROI, and used it to create a new 4mm subject-specific ROI used to extract the mean BOLD time-series signal. The mean time series of the seed ROI and the contrast of interest (i.e. food cues vs. scenery images) were multiplied to create the PPI regressor in the first level analysis. Higher-level analysis was repeated as stated above.

For all of the univariate results reported here, the significance of the clusters was determined by threshold-free cluster enhancement (TFCE) with a significance level of p<0.05 family wise error rate (FWER). All coordinates are given in MNI space.

